# Hyperiid amphipods from the Gulf of Ulloa and offshore region, Baja California: Intermittent use of the coastal shelf

**DOI:** 10.1101/2020.04.29.068007

**Authors:** Bertha E. Lavaniegos

## Abstract

Hyperiid amphipod species from the Gulf of Ulloa, Baja California, and the adjacent region (from shelf break to 200 km offshore) were analyzed in order to evaluate diversity and abundances in this productive area that supports small-scale commercial fisheries such as barred sand bass (*Paralabrax nebulifer*), California spiny lobster (*Panulirus interruptus*), abalones, clams, and others. Strong coastal upwelling events were observed during summer seasons of the period 2002-2008 between Punta Eugenia and Punta Abreojos. The upwelling plumes at Punta Abreojos are projected southward in slope waters bordering the coastal shelf of the Gulf of Ulloa, contributing to the separation of coastal and oceanic regions, and explain differences in amphipod diversity and abundances. In the offshore region, the most abundant species were *Vibilia armata, Lestrigonus schizogeneios, Primno brevidens*, and *Eupronoe minuta*, similar to previous findings in northern regions of Baja California and southern California. However, their abundances were lower (between 10 and 30 individuals/1000 m^3^), only reaching 20-50% of abundance levels reported off northern Baja California. In the coastal shelf of the Gulf of Ulloa, amphipods were virtually absent during 2002, 2003 and 2006. However, during 2004 and 2005, abundances of *P. brevidens* increased (54 and 20 ind/1000 m^3^, respectively). Moreover, during 2007, abundances of *L. schizogeneios, P. brevidens, Lycaea nasuta, Lycaea pulex*, and *Simorhynchotus antennarius* increased considerably (261, 39, 31, 68, 416 ind/1000 m^3^, respectively), indicating occasional utilization of the coastal shelf by pelagic amphipods. Gelatinous organisms paralleled changes in hyperiid populations and were particularly abundant in 2007 in the coastal shelf. Significant correlations of 17 amphipod species with gelatinous taxa, which are often used as host organisms by hyperiid amphipods, suggest that those organisms enhanced hyperiid abundance and promoted the progression onto the coastal shelf during some years of the 2002-2008 period.

## Introduction

The zooplankton community has been intensively studied in the northern regions of the California Current System (CCS), but off Baja California it has received less attention, particularly in terms of taxa such as the hyperiid amphipods that comprise only a small proportion of the community and are therefore assumed to have minimal ecological importance. However, this perception may be incorrect for oceanic waters, where they are relatively abundant [1-4] and may represent attractive forage food for predators [5-10]. Hyperiid amphipods have maximum abundances in high latitudes [11,12], while in tropical and subtropical regions they have low abundance but higher species diversity [13,14]. However, information from wide areas of tropical and subtropical seas is sparse and more studies are required in order to have a more precise quantification of these crustaceans and their role in trophic webs.

Seasonal changes in hyperiid amphipod species assemblages have shown a strong coupling with upwelling dynamics off Oregon [15], and the southern sector of the CCS, is to say, southern California [16] and Baja California [1-2]. Lowest abundances occur in winter, followed by increases in spring-summer, and maximum levels in autumn. However, hyperiids appear to avoid the zone of highest upwelling activity, as suggested by their relative scarcity in inshore waters compared to higher concentrations in the core and offshore part of the California Current [1,15]. The oceanic distribution of hyperiids, with mean abundance by species on the order of <1 to 100 ind/1000 m^3^ contrast with zero to 40 ind/1000 m^3^ in the coastal shelf [1]. The difference is more remarkable considering to the main species responsible of the secondary production at Vizcaino Bay, the copepods *Calanus pacificus* and *Acartia tonsa*, which reach 138 and 60 ind m^−3^, respectively, on the coastal shelf [17], and the euphausiid *Nyctiphanes simplex* with 35-87 ind m^−3^ [18]. The high abundance of *C. pacificus, A. tonsa* and *N. simplex*, in Vizcaino Bay and the Gulf of Ulloa [17,19] explains the attraction of fishes and large predators to these neritic waters. Mesoscale structures, particularly eddies, could contribute to concentrate the high biological productivity in the Gulf of Ulloa [20]. However, the region also presents seasonal changes in circulation, with a poleward flow from July to October nearshore [21], which could bring oceanic fauna onto the shelf. Therefore, the region appears to have a dynamic circulation promoting high productivity that supports the small-scale coastal fishery at Punta Abreojos [22,23] and makes it a hotspot for tuna [24], swordfish [25], whales [26], and sea turtles [27,28].

In order to expand our understanding of zooplankton community and mechanisms influencing their variability in the southern part of the CCS, the present research describes interannual variability in summer species composition of hyperiid amphipods during 2002-2008. I will present evidence of intermittent occupation of the coastal shelf in the Gulf of Ulloa, Baja California, by hyperiids, and possible mechanisms underlying this behavior.

## Materials and methods

The study area is located off southwest Baja California between Punta Eugenia and Cabo San Lazaro and is characterized by a relatively wide coastal shelf (Fig 1). The coastline maintains a NW-SE orientation, favorable to upwelling [29], reinforced by the narrow step-shelf between Punta Eugenia and Punta Abreojos. Further south, the coastal shelf broadens to form the embayment named the Gulf of Ulloa (GU). Upwelling events induced by trade winds (Easterlies) are particularly frequent during spring and summer [30]. The lowest sea surface temperature (SST) occurs in February-April (18°C) while the maximum is recorded in summer (25°C), associated with an enhanced poleward current [21]. Poleward flow produces a decrease in chlorophyll, contrasting with the high productivity during spring (250-750 mg C m^−2^ d^−2^), when equatorward flow and strong upwelling is dominant [20].

**Fig 1.**
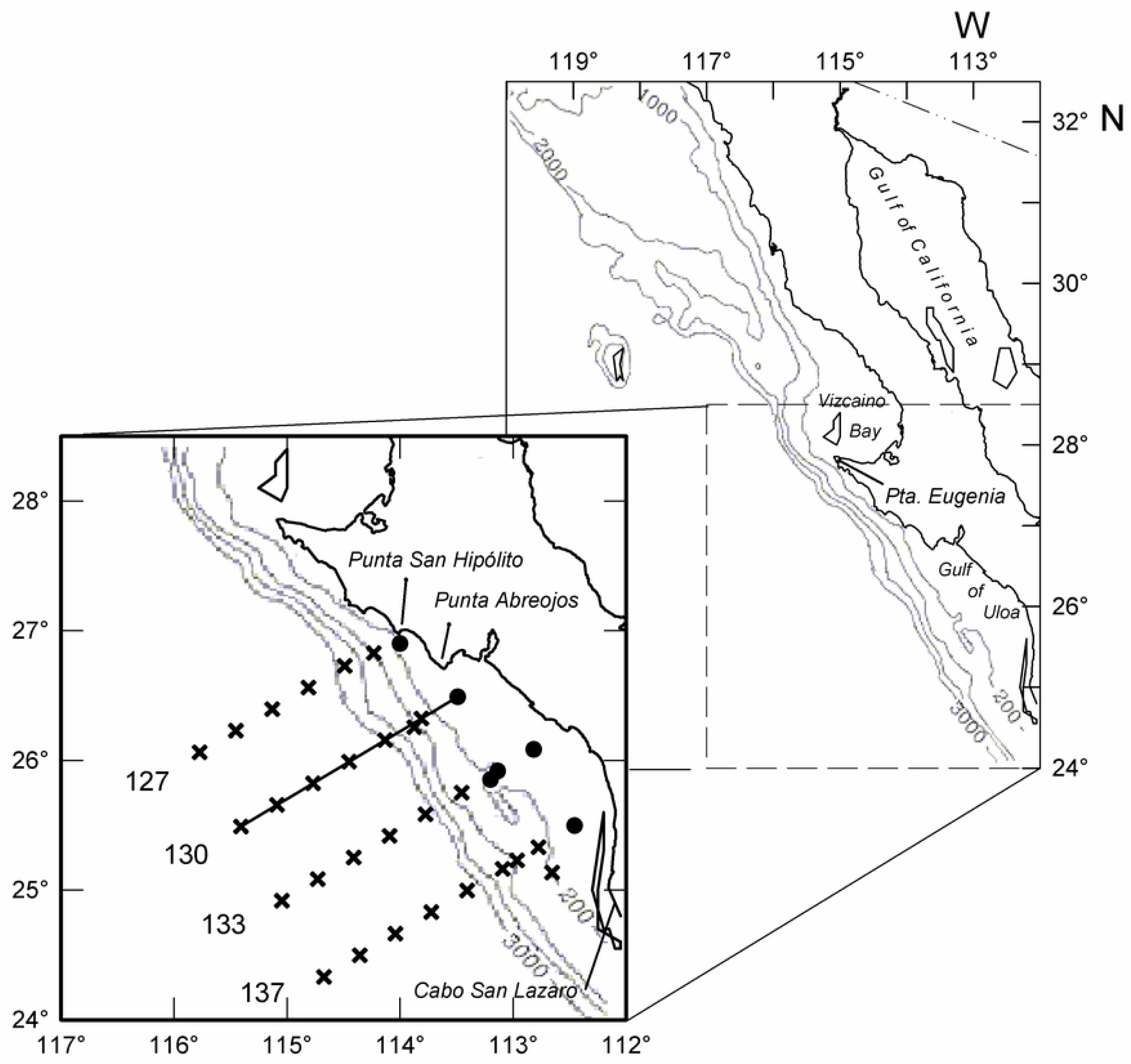
Study area. Gulf of Ulloa and offshore region showing the sampling grid and bathymetry (m). Stations analyzed in each cruise was variable (see S1 Table). The 200 m isobath separates coastal and oceanic stations (round and cross symbols respectively). Solid line indicates the transect used in vertical profiles. Numbers are transect-lines.

The sampling stations were arranged in four transect-lines perpendicular to the coast (Fig 1) from seven cruises performed every summer from 2002-2008 by the IMECOCAL program (Spanish acronym of: *Mexican Investigations of the California Current*). At each station, hydrographic data were recorded with a CTD (Seabird Electronics Inc., 9/11). Zooplankton sampling was carried out by oblique tows of a bongo net (71 cm-diameter, 505 µm mesh width) with a digital mechanical flowmeter. In the epipelagic oceanic region, tows were conducted from 0-210 m, while at stations located in the coastal shelf the tow depth was from the surface to 10-15 m above to the bottom. The samples were preserved with 4% formalin.

Only the nighttime samples were selected to perform taxonomic analysis of hyperiid species to reduce variability due to vertical migration, as has been reported for some hyperiid species (p. ej. [31,32]. However, for the coastal shelf (bottom depth < 200 m) daytime samples were also included due to the low number of stations. The cruise dates and number of samples analyzed are shown in Table 1 (details of sampling stations in S1 Table). Species identification was based on taxonomic keys [13,33]. Specimens of five gelatinous groups (medusae, siphonophores, ctenophores, doliolids, salps) were also counted, given their important role as host of amphipods [34,35].

**Table 1.** IMECOCAL cruises with dates and number of zooplankton samples used in taxonomic identification. Cruise dates correspond to the area considered in the present study (see Fig 1). The sampling hour is indicated as daytime (D) and nighttime (N).

| Cruise | Date | Number of samples |  |  |
| --- | --- | --- | --- | --- |
|  |  | Offshore region | Gulf of Ulloa |  |
|  |  | N | D | N |
| 0207 | 26 Jul–1 Aug 2002 | 10 | 2 | 3 |
| 0307 | 23–27 Jul 2003 | 8 | 3 | 1 |
| 0407 | 23–29 Jul 2004 | 8 | 3 | 3 |
| 0507 | 29 Jul–4 Aug 2005 | 10 | 4 | 2 |
| 0607 | 19–24 Jul 2006 | 8 | 1 | 4 |
| 0708 | 6–13 Sep 2007 | 12 | 4 | 2 |
| 0807 | 14–21 Jul 2008 | 14 | 3 | 3 |

In the present study, species occurring in 31-80% of samples will be considered dominant, those present in 10-30% of samples as common, and species with <10% of positive records as rare species. Analysis of variance (ANOVA) was applied to compare abundances among years for the most abundant species and gelatinous zooplankton groups using log-transformed data (log_10_ [x-+1]). Further, *a posteriori* comparison with the Tukey test was applied to assess the specific years with differences. Transformed data were also used to address multivariate community analysis based in the Euclidian distance, and clusters were defined with the Ward’s linkage hierarchical method [36]. After exclusion of 16 very rare species (occurring in only one sample) as well as 7 samples from coastal stations without amphipods, the data matrix was of 75 species × 101 samples. Analysis of similarities (ANOSIM) was done to test the hypothesis for differences between clusters based in a resemblance matrix of Bray-Curtis index [37].

The symbiotic relation between hyperiid species and gelatinous zooplankton groups was analyzed using Spearman correlation analysis.

## Results

### Physical environment

Vertical sections of temperature in Line 130, extending from Punta Abreojos to 250 km offshore (Fig 2a-g), showed strong stratification during summer. Sea surface temperatures were slightly lower in 2002, 2005, and 2008, ranging from 19-22°C offshore and 18-20°C inshore. Maximum SST values were recorded in 2007 (22-23°C) because sampling was conducted in late summer (Fig 2f) when SST increases compared to mid-summer, the period when all other years were sampled. The upwelling footprint throughout the study period was more evident in Station 130.35 offshore than in the most coastal Station 130.30. Despite low SST at Station 130.35, temperature at 100-200 m depth was consistently higher than the rest of stations in Line 130 due to the influence of the California Undercurrent.

**Fig 2.**
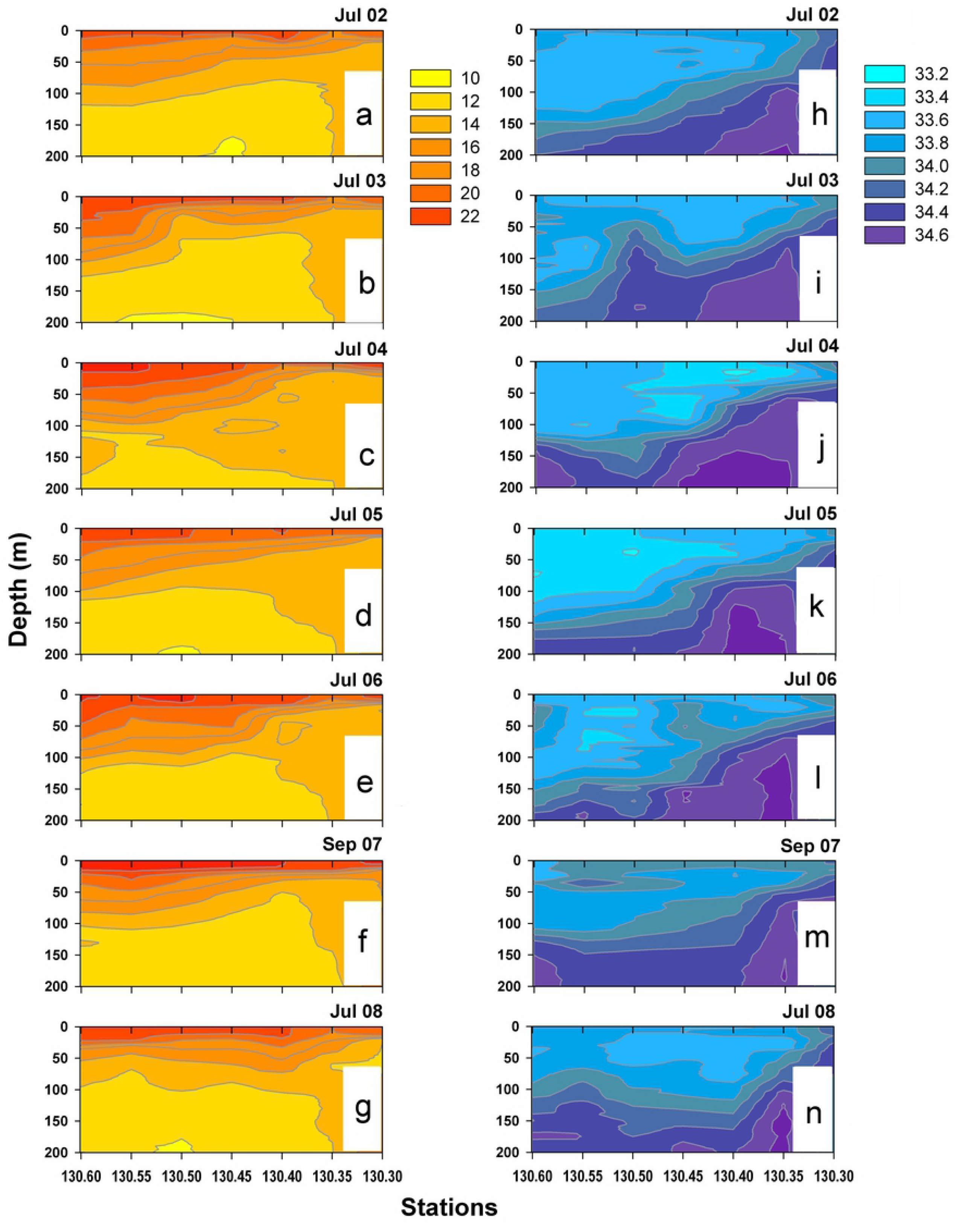
Vertical profiles. Temperature (a-g) and salinity (h-n) in the upper 200 m along the line 130 during the summers of 2002-2008.

A high salinity core in station 130.35 indicates the influence of the California Undercurrent (Fig 2h-n). The tilting of the isohalines is pronounced toward the coast, indicating intense upwelling activity. In July 2002 and again in 2007-2008, high-salinity upwelled water (>34 psu) reached the coastal shelf, but from 2003-2006 low salinity water in the upper layer masked the upwelling.

In summary, thermohaline conditions off southern Baja California showed marked onshore-offshore differences during every summer season and evidence for intense upwelling activity in the slope region (Stn. 130.35). The main interannual differences were slightly lower SST in 2002, 2005, and 2008 compared to other summers, and the influence of low salinity water in 2002-2006.

### Distribution and abundance of hyperiid amphipods

Abundance of total hyperiid amphipods varied between stations from 0 to 3,732 ind/1000 m^3^. Except for 2007, hyperiids collected at GU had low abundance (< 150 ind/1000 m^3^) and were absent in seven stations (3 from 2002, 2 in 2003, 1 in 2005, and 1 in 2008; Fig 3). Approximately 37% of GU samples had abundances higher than 170 ind/1000 m^3^ in 2004, 2005, and 2007, and four stations in 2007 surpassed 1000 ind/1000 m^3^. Thus, excepting summer 2007, amphipods showed a clearly oceanic tendency, indicated by higher presence and abundance in offshore stations with bottom depth >200 m. In the offshore region, the highest abundances were observed in 2007 and 2008, and lowest abundance in July 2003 (Fig 3).

**Fig 3.**
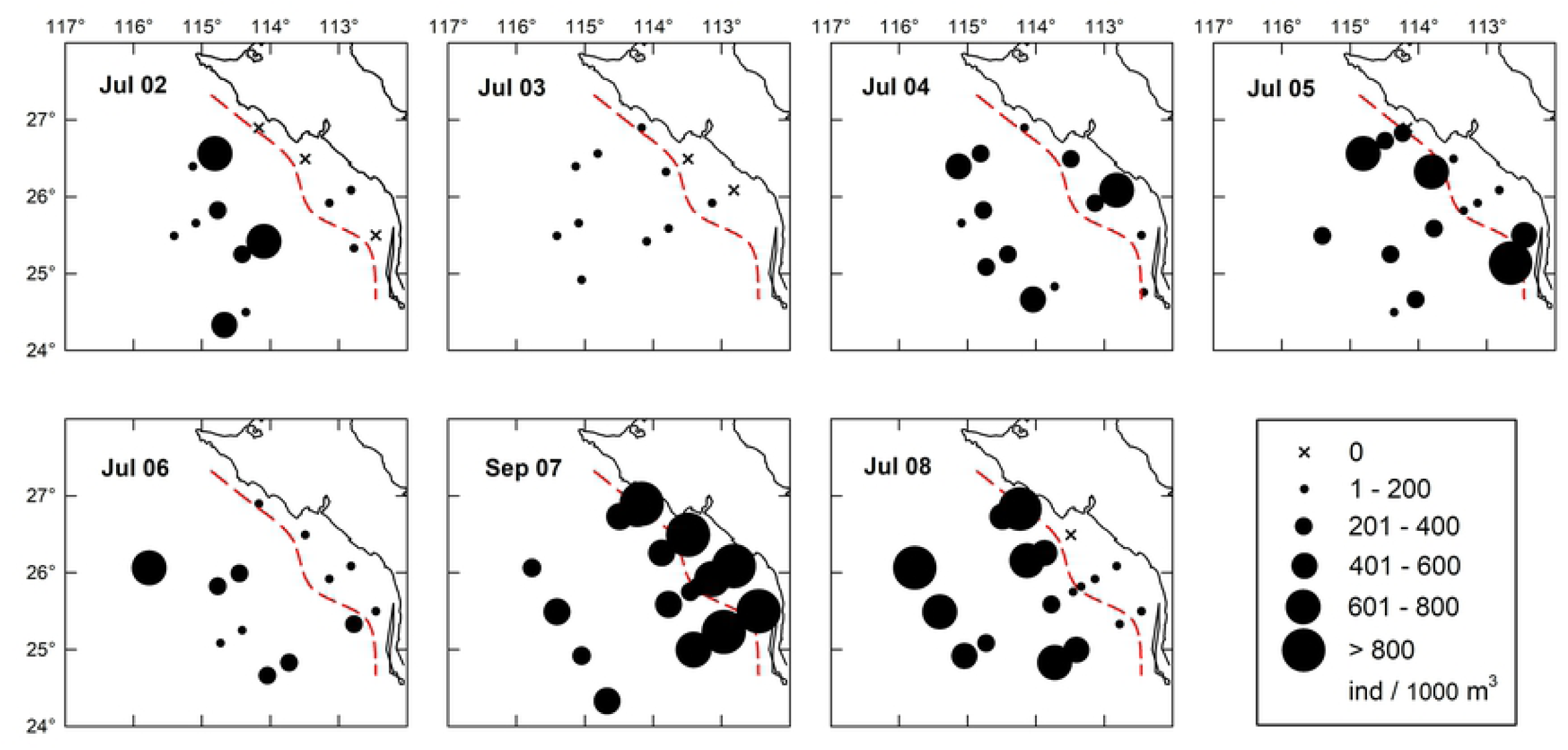
Distribution of hyperiid amphipods. Total abundance distribution in the Gulf of Ulloa and offshore region during the summers of 2002-2008. Red dashed line divides the onshore and offshore stations.

The ANOVA comparing abundances across years in the offshore region was significant, and summer 2003 was notably lower than all other years (F = 11.2, p < 0.001). The geometric mean (GM) in 2003 was 48 ind/1000 m^3^, and it fluctuated between 190 and 445 ind/1000 m^3^ in the other years (Fig 4a). In the GU, hyperiid abundance also presented significant differences (F = 9.1, p < 0.001). The highest abundance recorded in 2007 in the GU (GM = 1,316 ind/1000 m^3^) was significantly higher than for 2002, 2003, 2006, and 2008, when the GM ranged from 2 to 27 ind/1000 m^3^ (Fig 4b). The abundance in 2004 (GM = 129 ind/1000 m^3^) was also significantly higher in relation to 2002 and 2003 (p = 0.020 and 0.010, respectively); 2005 (GM = 78 ind/1000 m^3^) had significantly higher abundance only compared to 2003 (p = 0.036).

**Fig 4.**
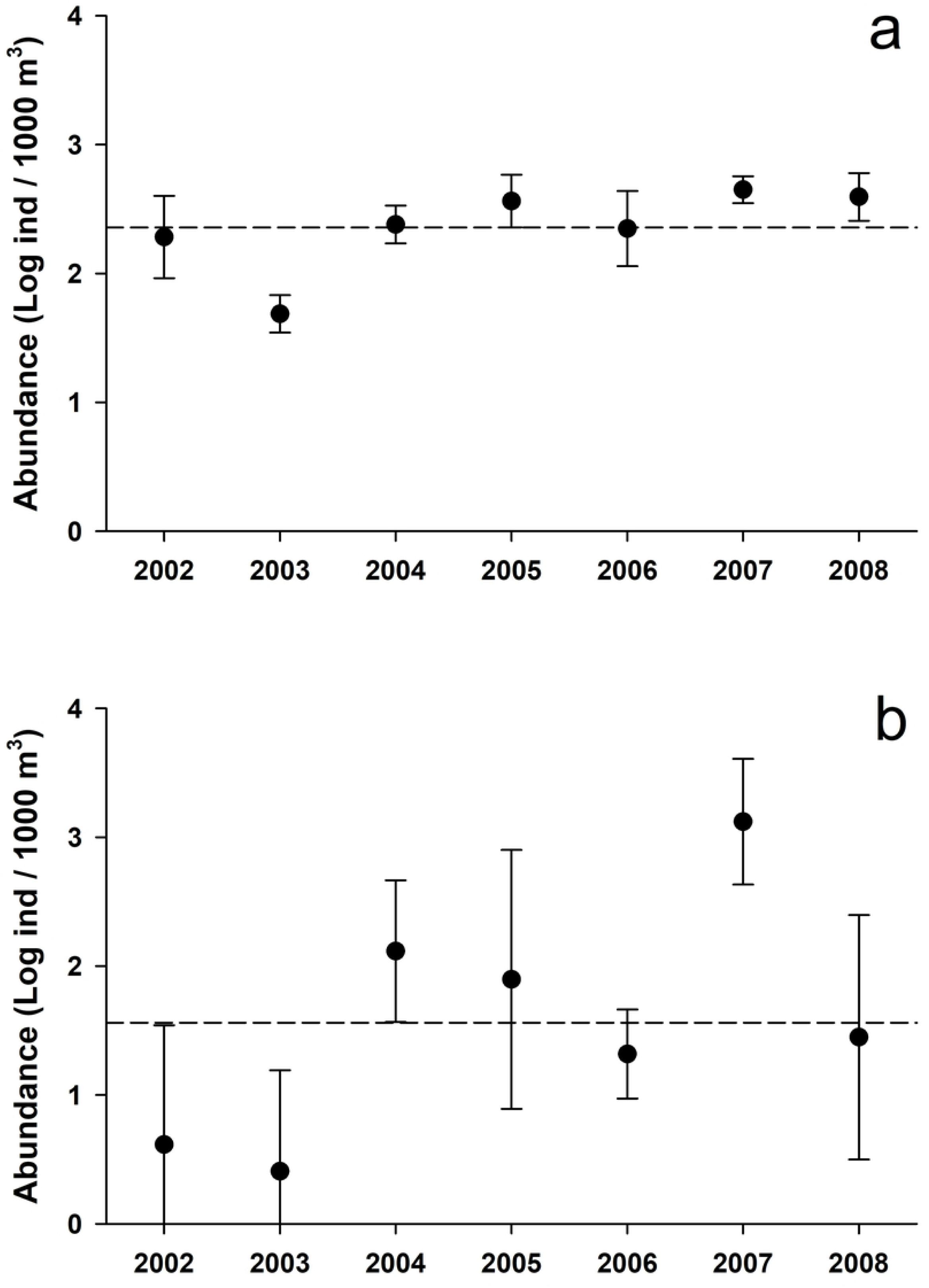
Summertime abundance of hyperiid amphipods. Mean (± 95% confidence interval) of total hyperiids in the oceanic region off south Baja California (a) and the Gulf of Ulloa (b), with data transformed to logarithms. The long-term mean is indicated by the dashed line.

#### Interannual variability of hyperiid species

Hyperiid amphipod diversity was high in the study region: 91 species were recorded in the 2002-2008 period (S2 Table). Eighty-eight of these species occurred in the oceanic region, while 56 were in neritic waters. There were 17 dominant species in the whole region, which appeared with a frequency of 31-80%, and had high abundances (GM between 1 and 15 ind/1000 m^3^). Only one dominant species pertained to the infraorder Physosomata (*Scina tullbergi*) while the rest were from the infraorder Physocephalata, with the main component in the superfamily Phronimoidea (9 species), followed by the superfamily Platysceloidea (5 species), and finally the superfamily Vibiloidea (2 species: *Paraphronima gracilis* and *Vibilia armata*).

Common species occurred with moderate frequency (10-30%) and GMs in the range of 0.1 – 0.9 ind/1000 m^3^. They included one species from Physosomata (*Scina borealis*) and 32 from Physocephalata, spread over the superfamilies Vibiloidea (12%), Phronimoidea (28%), and Platysceloidea (59%). The complete list of species is shown in S2 Table. Almost all common species were missing in one or more years, but a few occurred across the period 2002-2008 (*Scina borealis, Lycaeopsis themistoides, Pronoe capito*, and *Rhabdosoma whitei*).

A high number of rare species presented low frequency (<10%), 7 from Physosomata and 36 from Physocephalata. Their global GM was below 0.1 ind/1000 m^3^ (S2 Table). The low presence of some rare species is probably due to their meso- and bathypelagic distributions, meaning they are rarely captured in the upper layer. Such is the case for *Lanceola clausi, Scypholanceola aestiva*, and *Scina curvidactyla* [13]. Most of the rare species only occurred in a single year (51%) and some occurred in only a single sample (42%). The year with the lowest number of rare species was 2003, and the highest was 2007, with 5 and 51% respectively. The rest of the years had 23-35%. Most of the rare species were found in the offshore region (72%); only 7% occurred exclusively inside the GU, and the remaining 21% occurred in both regions.

Interannual abundance comparisons within each dominant or common species using ANOVA were significant for 26 species in the offshore region, representing 76% of dominant species and 42% of common species (p<0.01, Table 2). A recurrent pattern was the high abundance during 2007, significantly higher than all other years, depicted for *Simorhynchotus antennarius, Vibilia stebbingi, Lycaea nasuta*, and *Oxycephalus clausi* (Fig 5a). In other species as *S. tullbergi* and *Hyperoche medusarum*, the abundance was similarly high in 2007 and 2008 compared to 2006. In *Laxohyperia vespuliformes* the abundance was high in 2007 and 2005 compared to 2002-2004 (see S1-S3 Figs).

**Table 2.** Interannual comparisons of hyperiid species. Abundances of dominant and common species compared with ANOVA. Only species with significant results are shown (α < 0.01). Specific years with significant differences are indicated, which resulted from Tukey multiple comparisons.

| Species | F | p | Multiple comparisons |
| --- | --- | --- | --- |
| **Offshore region (N = 70)** |  |  |  |
| *Amphithyrus sculpturatus* | 8.2 | <0.001 | (2007,2008) < (2002,2004,2005)<br>2003 < 2002 |
| *Anchylomera blossevillei* | 4.4 | <0.001 | (2003,2005,2006) < 2004 |
| *Eupronoe armata* | 4.2 | 0.001 | (2002,2003,2005,2008) < 2007 |
| *Eupronoe maculata* | 4.4 | <0.001 | (2003,2004,2005) < 2002 |
| *Eupronoe minuta* | 5.7 | <0.001 | 2003 < other years |
| *Hyperioides longipes* | 7.4 | <0.001 | other years < 2005 |
| *Hyperioides sibaginis* | 3.8 | 0.003 | (2002,2005) < 2004 |
| *Hyperoche medusarum* | 3.6 | 0.004 | 2006 < (2007,2008) |
| *Laxohyperia vespuliformis* | 6.8 | <0.001 | (2002,2003,2004) < (2005,2007)<br>2006 < 2007 |
| *Lestrigonus bengalensis* | 3.5 | 0.005 | 2002 < (2004,2005,2008) |
| *Lestrigonus macrophthalmus* | 5.2 | <0.001 | (2002,2003,2005,2007,2008) < 2004<br>(2005,2007) < 2006 |
| *Lestrigonus schizogeneios* | 6.4 | <0.001 | 2003 < (2002,2004,2005,2007,2008) |
| *Lycaea nasuta* | 5.3 | <0.001 | other years < 2007 |
| *Lycaeopsis zamboangae* | 11.1 | <0.001 | other years < 2004<br>(2002,2005) < 2007 |
| *Oxycephalus clausi* | 6.9 | <0.001 | other years < 2007 |
| *Phronima curvipes* | 4.1 | 0.002 | (2003,2004,2007) < 2008 |
| *Phronima sedentaria* | 3.4 | 0.006 | (2005,2007) < 2008 |
| *Phrosina semilunata* | 4.4 | <0.001 | 2007 < (2005,2006,2008)<br>2003 < 2005 |
| *Platyscelus ovoides* | 4.2 | 0.001 | (2003,2004) < 2002 |
| *Primno brevidens* | 5.9 | <0.001 | 2003 < (2005,2006,2008) |
| *Scina tullbergi* | 12 | <0.001 | (2002,2003,2004,2006) < (2007,2008) |
| *Simorhynchotus antennarius* | 17.9 | <0.001 | other years < 2007<br>(2002,2003,2005) < 2008 |
| *Themistella fusca* | 7.9 | <0.001 | (2002,2003,2005,2008) < (2004,2007) |
| *Tryphana malmi* | 4.3 | 0.001 | other years < 2008 |
| *Vibilia stebbingi* | 10.7 | <0.001 | other years < 2007 |
| *Vibilia viatrix* | 6.1 | <0.001 | (2003,2005,2006,2007,2008) < 2002 |
| **Coastal shelf (N = 38)** |  |  |  |
| *Eupronoe minuta* | 5.8 | <0.001 | (2002,2003,2006) < 2007 |
| *Laxohyperia vespuliformis* | 4.6 | 0.002 | (2002,2004,2006) < 2007 |
|  |  |  | 2004 < 2005 |
| *Lestrigonus schizogeneios* | 10.7 | <0.001 | other years < 2007 |
| *Lycaea nasuta* | 15.5 | <0.001 | other years < 2007 |
| *Lycaea pulex* | 13.0 | <0.001 | other years < 2007 |
| *Phronima atlantica* | 9.4 | <0.001 | other years < 2007 |
| *Platyscelus ovoides* | 3.7 | 0.007 | (2002,2003,2004,2006,2008) < 2007 |
| *Rhabdosoma whitei* | 3.5 | 0.009 | other years < 2007 |
| *Simorhynchotus antennarius* | 35.8 | <0.001 | other years < 2007 |
|  |  |  | (2002,2003,2004,2006) < 2008 |
| *Tryphana malmi* | 6.5 | <0.001 | (2002,2003,2004,2005,2006) < 2007 |
| *Vibilia stebbingi* | 11.9 | <0.001 | other years < 2007 |

**Fig 5.**
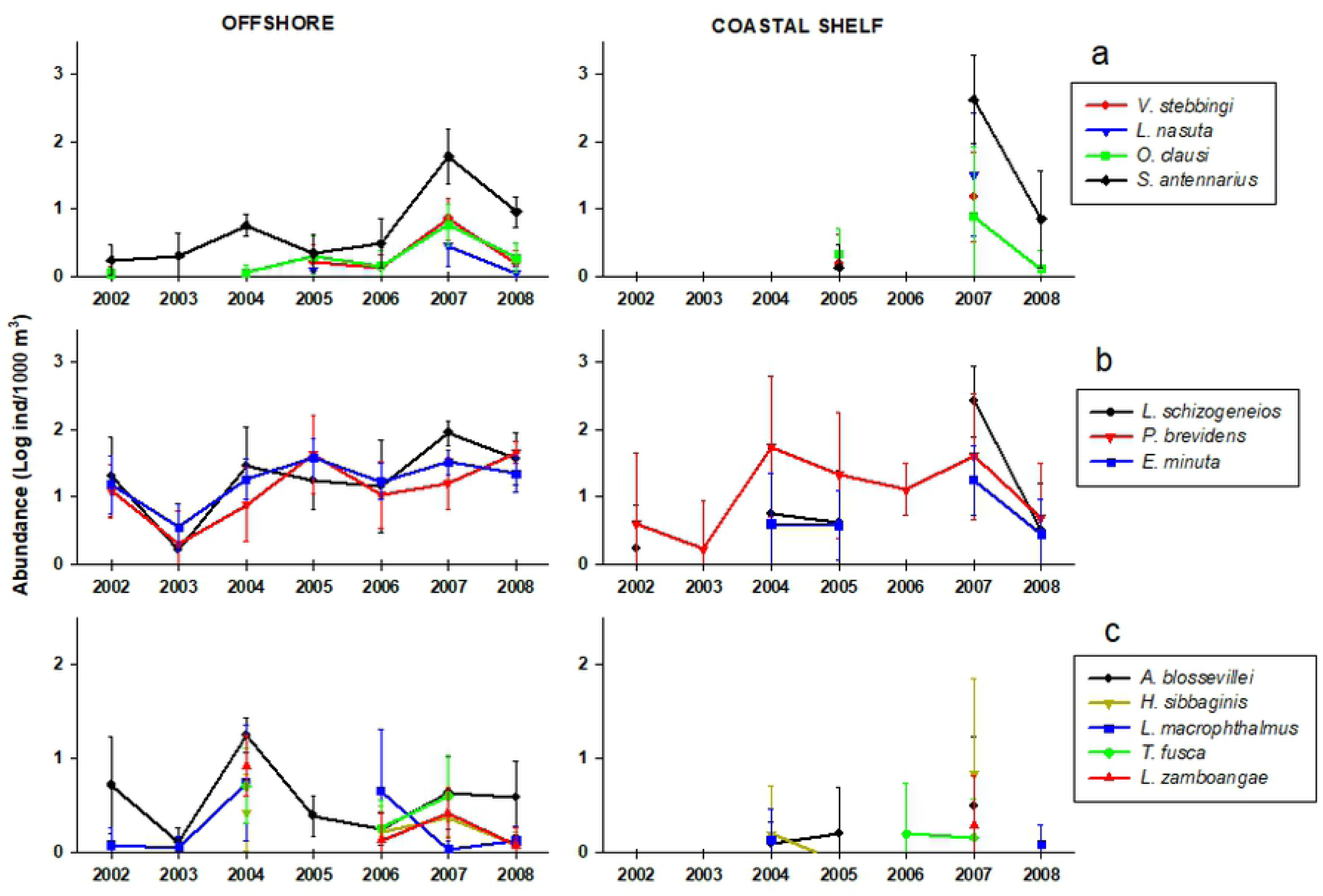
Patterns of interannual variability. Mean (± 95% confidence interval) in species groups showing an increase in 2007 (a), decreasing in 2003 (b), or increasing in 2004 (c).

In general, 2007 and 2008, and to a lesser extent 2004, appeared to be years of high abundance for many species, while 2002, 2003, and 2006 were years of low abundance. Some species showed a uniform pattern throughout the entire study period except for a strong decrease in 2003, particularly remarkable in the dominant species *Lestrigonus schizogeneios, Primno brevidens*, and *Eupronoe minuta* (Fig 5b). Summer 2004 was notable for high abundances *of Anchylomera blossevillei, Hyperioides sibbaginis, Lestrigonus macrophthalmus, T. fusca*, and *Lycaeopsis zamboangae* (Fig 5c).

In the coastal shelf region, 12 species showed significant interannual differences (p<0.01, Table 2). In all cases, abundances were the highest in 2007 and generally lowest in 2002, 2003, and 2006. During 2007, *Lestrigonus bengalensis, L. schizogeneios*, and *S. antennarius* reached higher GM in the coastal shelf (28, 261, and 416 ind/1000 m^3^) compared to the offshore region (3, 87, and 59 ind/1000 m^3^; Fig. 5, S2-S3 Figs).

#### Assemblages of hyperiid species

Multivariate analysis resulted in four main clusters with variable number of stations (indicated with letters A-D in Fig 6). Cluster A may be split into two subgroups, one mainly oceanic (A1) and other neritic (A2). The oceanic stations in the A1 assemblage were from 2003, and one coastal station from 2004 (137.25) was joined to this cluster. Assemblage A2 contained two oceanic stations from 2002 (137.30, 137.55) but the rest were coastal stations from mixed years excluding 2005 and 2007 (Fig 7). Cluster B was exclusive for 2007, grouping five coastal shelf stations and four stations external but near the shelf (127.36, 137.33, 137.35, 137.40). The cluster C was also dominated by coastal stations and is divided into two subgroups of different years: C1, which consists of 14 stations (seven from 2008, six from 2007, and one from 2005), and C2, which has a mixture of stations from 2002-2006. The largest cluster, D, consisted of 44 oceanic stations, split in two subgroups: D1, combining eight samples from 2004 and three from 2007; and D2, including 33 samples from 2002, 2005, 2006, and 2008 (Figs 6 and 7).

**Fig 6.**
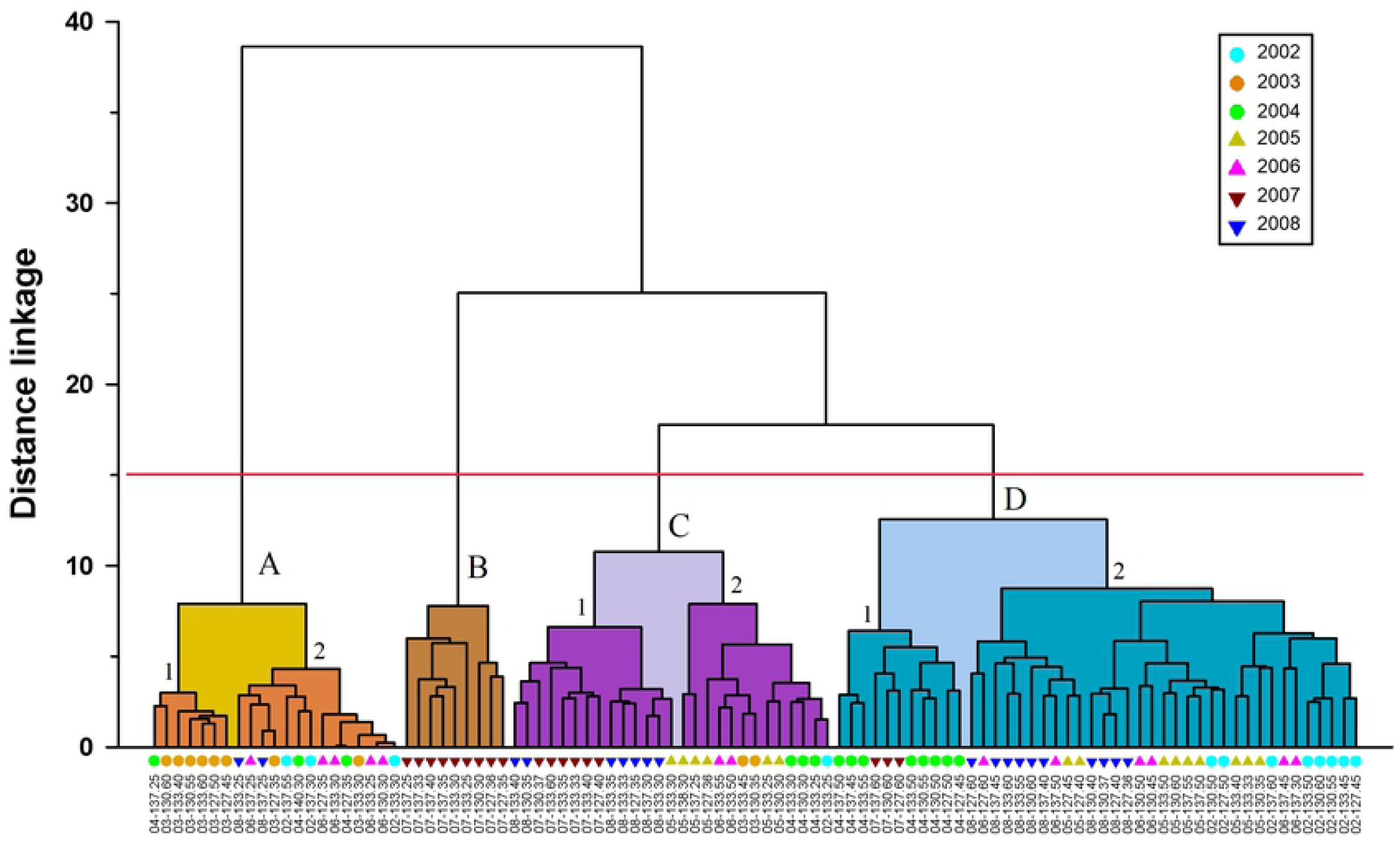
Hyperiid assemblages. Clustering of sampling stations based in abundances of 75 species. Clusters formed at a distance linkage of 25 (cutoff line in red) designed by letters (A, B, C, D), and subgroups by numbers. In the x-axis the sampling stations are shown with symbols in color indicating the year.

**Fig 7.**
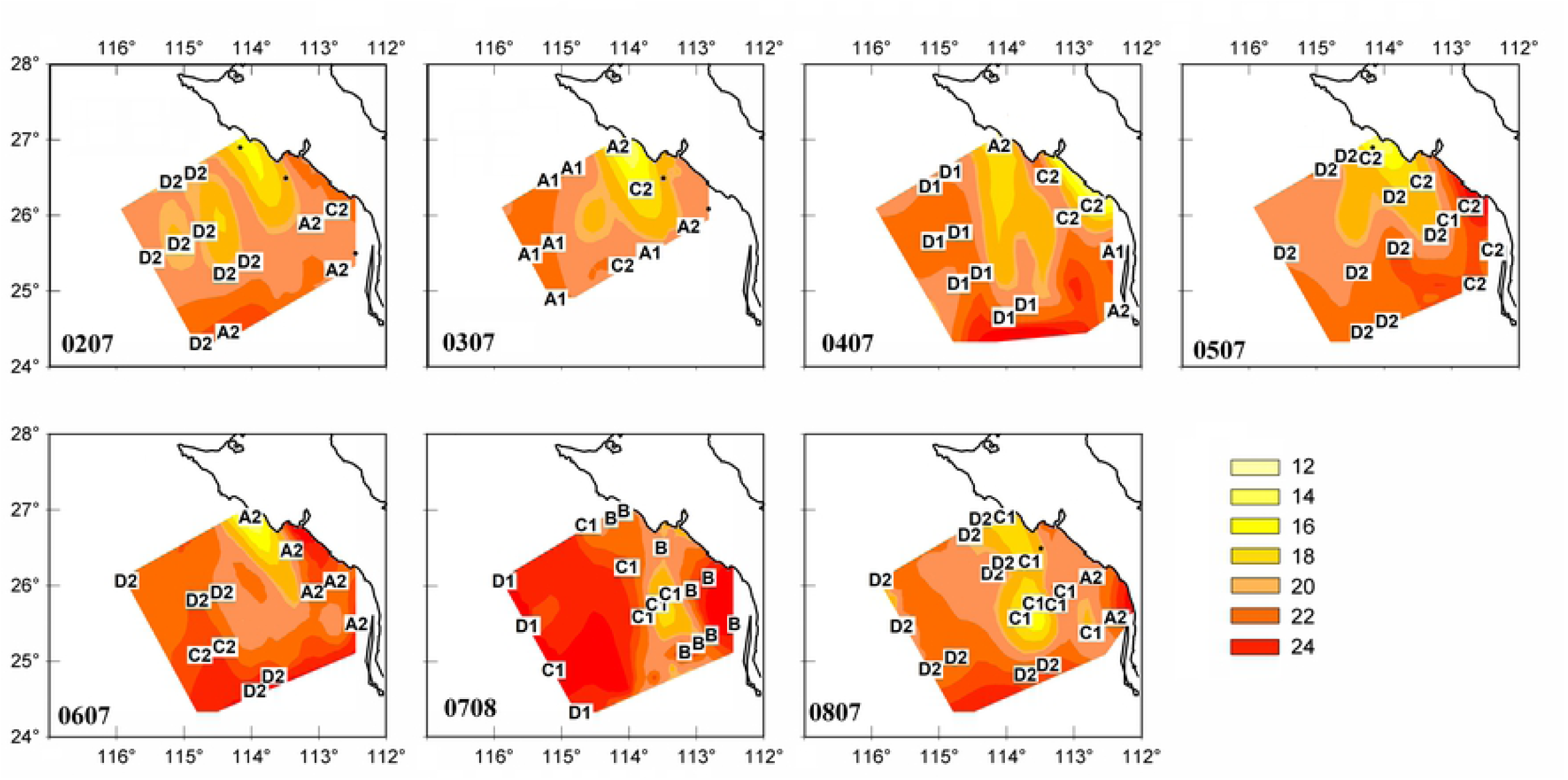
Geographic distribution of clusters. Clusters defined in Fig 6 are shown and sea surface contours (°C) were added as a reference to climatic conditions. Black points indicate stations without amphipods.

The differences among the seven clusters, including groups and subgroups, were confirmed with ANOSIM, which had a global R = 0.626 (p = 0.001). The pairwise tests between clusters showed also significant differences in all cases (p = 0.001).

The clusters considered coastal showed strong contrasts in species composition between the cluster B and the other three (A2, C1 and C2, Fig 8). Cluster B did not include any data from 2007 (Fig 7), and had high abundances of *S. antennarius, L. schizogeneios, Lycaea pulex*, and *P. brevidens*, with geometric means (GM) of 343, 246, 61 and 51 ind/1000 m^3^ respectively (Fig 8a). The Bray-Curtis similarity for these four species was the highest in cluster B, combining 44.6% (Fig 8b). The coastal cluster C1 included samples from 2017-2018 and was somewhat analogous to cluster B in the contribution to similarity by *S. antennarius, L. schizogeneios*, and *P. brevidens* (summing 41.9%) but with lower abundances. The main difference between clusters B and C1 was due to *Lycaea nasuta* and *L. pulex*, which contributed 8.8 and 5.5%, respectively, to the similarity of cluster B; cluster C1 contributed 0.2 and 0%.

**Fig 8.**
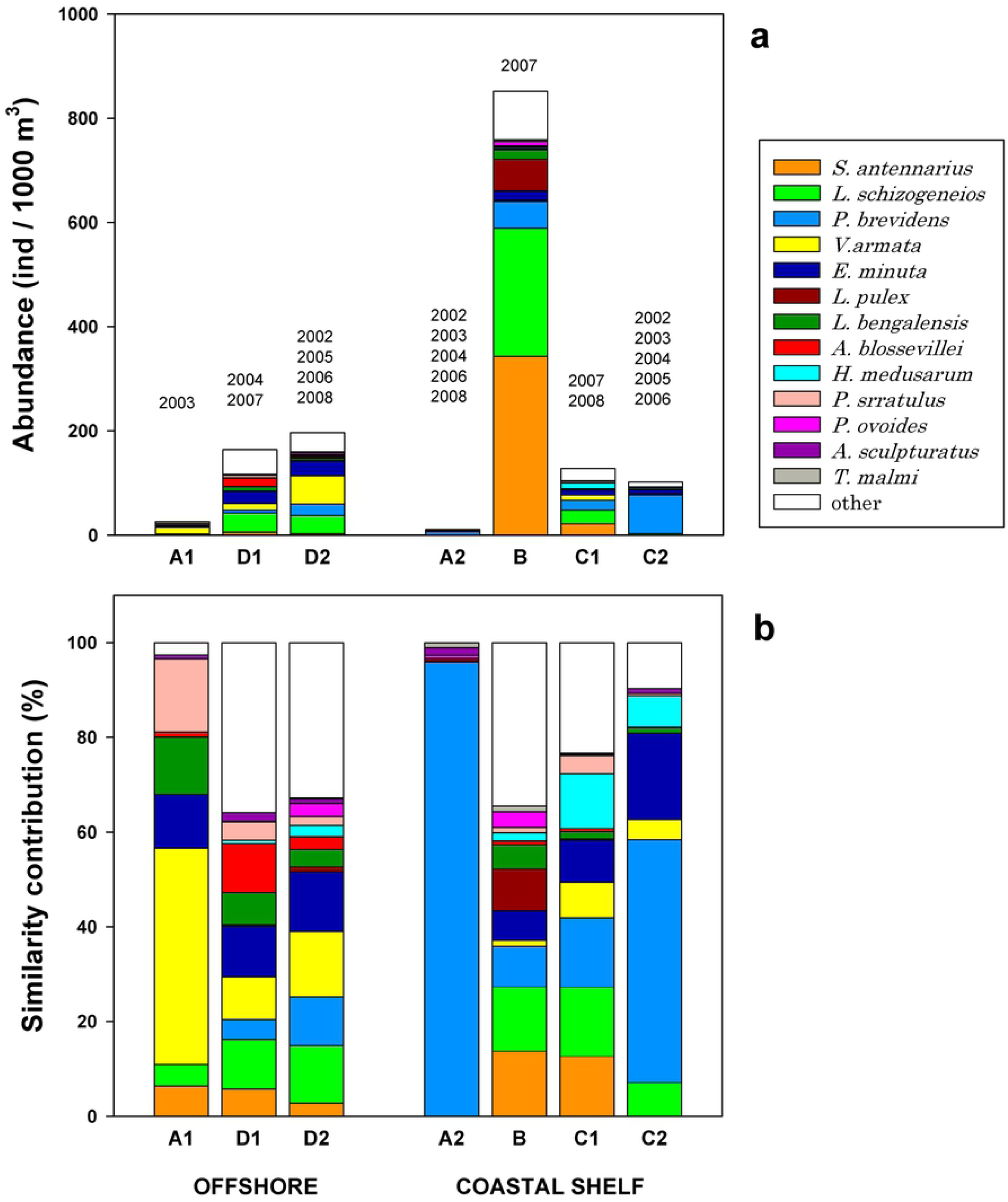
Species contribution in Clusters. Stacked geometric means (a) and contribution to similarity (b) of main species in clusters defined in Fig 6. The selected species are a combination of the four with higher similarity percentages from each cluster. The numbers indicate the years with one or more stations in the clusters

In contrast, coastal clusters C2 and A2 showed the highest contribution to similarity by *P. brevidens*, with 51.2 and 95.9% respectively (Fig 8b). However, the high percentage in cluster A2 was due to a low diversity (23 species with abundances <0.6 ind/1000 m^3^) but also low abundance of *P. brevidens* (GM = 6 ind/1000 m^3^) compared to other coastal clusters. Instead, the assemblage C2 had abundant *P. brevidens* (74 ind/1000 m^3^) and 47 other species present, though only nine (*E. minuta, L. schizogeneios, H. medusarum, Hyperioides longipes, V. armata, L. vespuliformes, S. borealis, Amphityrus sculpturatus*, and *P. sedentaria*) were moderately abundant (1-8 ind/1000 m^3^). The assemblages of hyperiid species in clusters A2 and C2 represent typical neritic communities since data included were from different years, even though C2 was more characteristic of 2005 (Fig 7).

In the offshore region, the interannual separation in different clusters was also clear but slightly different from the neritic region (Figs 6-8). The most different assemblage was cluster A1, characteristic of 2003, with the highest contribution to similarity by *V. armata* (45.6%), followed by *P. serratulus* (15.5%) and *L. bengalensis* (12.1%). These species had low abundances (GM of 12, 3 and 2 ind/1000 m^3^ respectively). *P. brevidens*, important in other clusters, was absent in cluster A1, and *L. schizogeneios* had low contribution to similarity (4.6%).

The offshore community from 2004 and 2007, represented in cluster D1, showed certain singularity, with main contribution of *E. minuta, L. schizogeneios, A. blossevillei*, and *V. armata* (summing 40.5% of similarity). Therefore, in 2007 there was strong contrast between oceanic (D1) and neritic (B) assemblages, mainly given by differences in *S. antenarius, L. pulex, V. stebbingi* and *L. nasuta*, which were scarce in cluster D1. The large cluster D2, characteristic of 2002, 2005, 2006, and 2008 (Fig. 7), had among the most representative species of *V. armata, E. minuta, L. schizogeneios*, and *P. brevidens* (accounting for 48.8% of similarity). As can be seen, cluster D2 was relatively similar to D1, with the main disparities being a higher abundance of *V. armata* (54 versus 13 ind/1000 m^3^) and *P. brevidens* (22 versus 5 ind/1000 m^3^), and lower abundance of *A. blossevillei* (3 versus 16 ind/1000 m^3^).

### Correlation of hyperiids and gelatinous plankton

Interannual changes of gelatinous organisms paralleled changes in hyperiid populations (Table 3). The low amphipod abundances observed during 2003 corresponded with low abundances of medusae, ctenophores, and doliolids in the offshore region, and virtually all gelatinous groups in the coastal shelf (Fig 9b). Low siphonophore abundance occurred in 2002 but remained at consistent levels from 2003-2008 in both regions (Fig 9). In the coastal shelf, siphonophore abundance changed by several orders of magnitude between 2002 (3 ind/1000 m^3^) and all other years in the time-period (GM ranging between 765 and 23,051 ind/1000 m^3^). In the offshore region, differences were moderate, with 421 ind/1000 m^3^ in 2002 compared to 1,992-6,130 ind/1000 m^3^ in other years.

**Table 3.** Interannual comparisons of gelatinous groups. Abundances of gelatinous groups compared with ANOVA. Taxa with significant results are highlighted (α < 0.01). Specific years with significant differences are indicated, which resulted from Tukey multiple comparisons.

| Taxa | F | p | Multiple comparisons |
| --- | --- | --- | --- |
| **Offshore region (N = 70)** |  |  |  |
| Medusae | 8.1 | **<0.001** | 2003 < other years |
| Siphonophora | 6.8 | **<0.001** | 2002 < (2004,2005,2006, 2007, 2008) |
| Ctenophora | 13.8 | **<0.001** | (2002, 2003,2006) < (2005,2007,2008) |
|  |  |  | 2004 < 2008 |
| Doliolida | 5.4 | **<0.001** | 2008 < (2004,2005,2007) |
|  |  |  | 2003 < 2007 |
| Salpida | 2.1 | 0.073 |  |
| All gelatinous organisms | 2.1 | 0.068 |  |
| **Coastal shelf (N = 38)** |  |  |  |
| Medusae | 4.5 | **0.002** | (2002,2003) < (2005, 2007) |
| Siphonophora | 11.0 | **<0.001** | 2002 < other years |
| Ctenophora | 2.1 | 0.077 |  |
| Doliolida | 4.8 | **0.002** | 2003 < (2005,2007) |
| Salpida | 3.9 | **0.006** | 2006 < 2008 |
| All gelatinous organisms | 2.7 | 0.034 |  |

**Fig 9.**
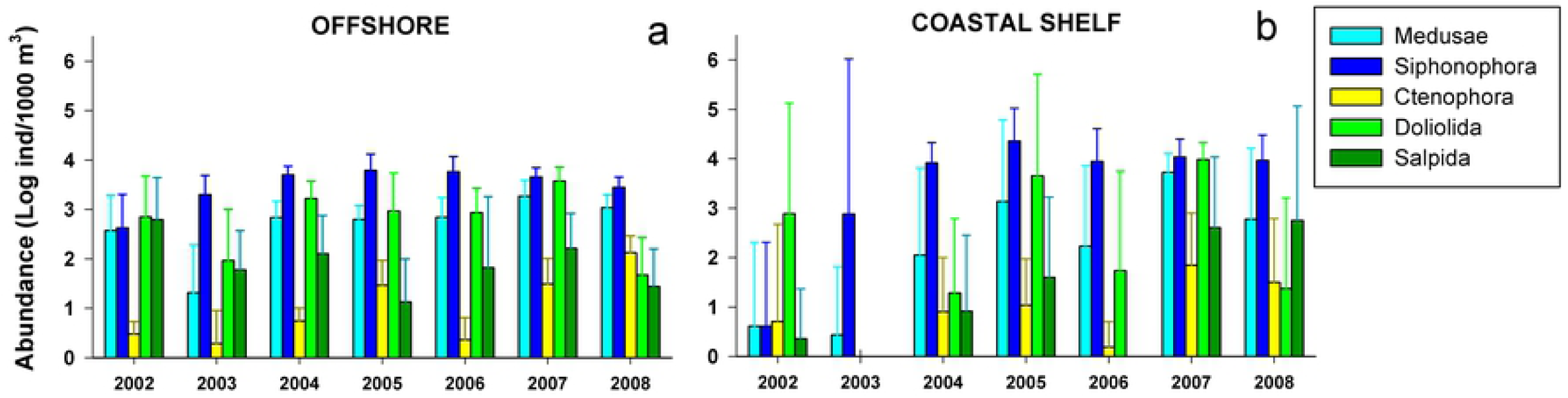
Gelatinous zooplankton. Mean (± 95% confidence interval) of gelatinous zooplankton groups in the offshore region (a) and coastal shelf (b).

Other gelatinous groups showed wide interannual variability and less coherence between regional patterns of variability. Medusae stand out in 2007 with particularly high abundance in the GU. Lowest abundance in both regions occurred in 2003 (GM of 20 and 2 ind/1000 m^3^ at offshore and GU, respectively; Fig 9, Table 3). In the offshore region, ctenophores were the least abundant gelatinous group throughout the study period but increased in 2005, 2007, and 2008 compared to 2002, 2003 and 2006 (Table 4). No significant differences were observed for ctenophores in the GU.

**Table 4.** Correlation between hyperiids and potential hosts. Spearman correlation between abundances of hyperiid amphipod species and gelatinous zooplankton groups. Significant correlations are highlighted (α < 0.001). Species without significant results are not shown.

| Hyperiid Species | Medusae | Siphonoph. | Ctenophora | Doliolida | Salpida |
| --- | --- | --- | --- | --- | --- |
| *Eupronoe maculata* | 0.228 | -0.103 | 0.133 | 0.143 | **0.405** |
| *Eupronoe minuta* | 0.174 | -0.023 | **0.389** | 0.225 | 0.193 |
| *Hyperioides longipes* | 0.217 | 0.216 | 0.084 | **0.398** | 0.096 |
| *Hyperoche medusarum* | **0.455** | 0.175 | **0.471** | 0.173 | 0.184 |
| *Laxohyperia vespuliformis* | **0.540** | **0.345** | **0.442** | **0.371** | **0.400** |
| *Lestrigonus schizogeneios* | **0.445** | 0.126 | **0.406** | **0.437** | 0.309 |
| *Lycaea nasuta* | **0.352** | 0.200 | 0.220 | **0.320** | 0.246 |
| *Lycaea pulex* | **0.344** | 0.126 | 0.159 | 0.264 | **0.342** |
| *Lycaea serrata* | 0.230 | -0.010 | 0.134 | 0.121 | **0.366** |
| *Oxycephalus clausi* | **0.409** | 0.268 | **0.376** | **0.341** | 0.310 |
| *Platyscelus ovoides* | **0.313** | 0.011 | 0.114 | **0.360** | **0.355** |
| *Platyscelus serratulus* | 0.245 | 0.125 | 0.288 | 0.172 | **0.322** |
| *Primno brevidens* | **0.383** | 0.285 | **0.457** | 0.179 | 0.019 |
| *Scina tullbergi* | 0.296 | 0.120 | **0.507** | 0.075 | 0.006 |
| *Simorhynchotus antennarius* | **0.455** | 0.209 | **0.430** | 0.285 | **0.355** |
| *Tryphana malmi* | 0.305 | 0.170 | **0.357** | 0.112 | 0.226 |
| *Vibilia stebbingi* | **0.479** | 0.222 | 0.305 | **0.480** | 0.292 |

Tunicates presented significant interannual differences at the GU but tendencies were different for doliolids and salps (Fig 9b, Table 3). Doliolids were absent in 2003 and their abundance in 2002 (GM = 772 ind/1000 m^3^) was significantly lower than in 2005 and 2007 (4,528 and 9,610 ind/1000 m^3^, respectively). Salps showed higher abundances in 2007-2008 (GM of 404-562 ind/1000 m^3^) but significant differences with the Tukey test were limited to 2006 and 2008 (p = 0.036). At the offshore region, doliolids had lower variability compared to the rough fluctuations observed in the GU. The pattern of salps was different between regions, with high abundance during 2002-2004 in the offshore region but low in the GU. In contrast, salps were more abundant in the GU during 2007-2008.

Correlation analysis between gelatinous groups and the 50 most frequent hyperiid species returned significant correlations for 17 species (p < 0.001, Table 4; 59% dominant and 41% common species). *L. vespuliformis* was exceptional in presenting significant correlation with all five gelatinous groups. Four other species (*L. schizogeneios, O. clausi, Platyscelus ovoides*, and *S. antennarius*) correlated positively with three gelatinous groups, but the rest of species only correlated with one or two gelatinous groups (Table 4).

The highest number of significant correlations (p<0.001) was with medusae and ctenophores (56% of the total). These included 4 species correlated with medusae (*V. stebbingi, L. nasuta, L. pulex*, and *P. ovoides*), 3 species with ctenophores (*S. tullbergi, E. minuta*, and *T. malmi*), and 7 species correlated with both medusae and ctenophores (*Hyperoche medusarum, L. vespuliformis, L. schizogeneios, P. brevidens, O. clausi*, and *S. antennarius*). There did not seem to be a taxonomic preference for one of the two groups, and all superfamilies had one or more species correlated with both medusae and ctenophores (Table 4).

Tunicates comprised 41% of significant correlations (p<0.001), with 5 species correlated with doliolids (*V. stebbingi, H. longipes, L. schizogeneios, L. nasuta*, and *O. clausi*), 5 correlated with salps (*Eupronoe maculata, L. pulex, L. serrata, Platyscelus serratulus*, and *S. antennarius*), and 2 species with both tunicate groups (*L. vespuliformis* and *P. ovoides*). All superfamilies excepting Scinoidea had some correlation with tunicates (Table 4). The siphonophores only presented one significant correlation, with *Laxohyperia vespuliformis*.

## Discussion

### Interannual variability and the weak El Niño events

The region located south of Punta Eugenia has the highest influence of tropical biota in the CCS, due to both resident and transient tropical species [1,38]. Despite high numbers of hyperiid amphipod species found in both the Gulf of Ulloa and the offshore region, abundances were low in the present study. Total hyperiids showed summer GM between 32-273 ind/1000 m^3^ during 2002-2008. In contrast, for the same period off northern Baja California (30-32°N), a GM in the range of 212-867 ind/1000 m^3^ was estimated [3]. The low amphipod abundance found in the GU could be related to El Niño events, which appeared to have a higher impact south of Punta Eugenia, based on changes in euphausiid populations [38]. At the species level, interannual changes in abundances of some species were out of phase between north-south Baja California regions. For example, the most abundant species, *Lestrigonus schizogeneios*, had a GM of <1 ind/1000 m^3^ in the present study, for the oceanic region during summer 2003, while off northern Baja California its GM was 75 ind/1000 m^3^ [3]. *Paraphronima gracilis* was the other species out of phase between northern and southern Baja California, with high abundance (GM = 77 ind/1000 m^3^) in the north region during 2003 [3], but low abundances off the GU (0.6 ind/1000 m^3^). The abundance of *P. gracilis* is in general low off the GU compared to northern Baja California (<3 ind/1000 m^3^ across the period 2002-2008). The low abundance during 2003 was also observed in eleven other species (Table 2).

It is interesting to note that the low hyperiid abundance recorded in 2003 south of Punta Eugenia was similar to the decrease that occurred during 2005 off northern Baja California [3]. In both cases, decreases were due largely to decreased abundance of *L. schizogeneios* and to a lesser extent *P. brevidens*. The question of but why these decreases occurred in different years from region to region may be answered by the occurrence of two short El Niño events in Jun 2002-Feb 2003 [39] and Jul 2004-Apr 2005 [40]. El Niño 2002-2003, though weak, could be related with the low hyperiid abundance in 2003 off the GU. The development of El Niño 2002-2003 was combined with a subarctic water intrusion coming from the north, which could have cooled the northern region off Baja California but not the southern region, due to the presence of a cyclonic eddy in the latitude of 27-29°N [41,42], helping to maintain the population densities of typical California Current species north of Punta Eugenia.

While El Niño 2002-2003 mainly affected southern Baja California, El Niño 2004-2005 appeared to more strongly affect hyperiid abundance in the northern region. Species such as *A. blossevillei* and *L. vespuliformis* had modest abundances in 2002-2008, but relatively more importance in the hyperiid community off the GU compared to the northern Baja California region [3]. *A. blossevillei* was particularly abundant in July 2004, comprising 11% of total hyperiids in that year. This species was also found during winter 2005 in the northern Baja California region [1], associated with oligotrophic waters, and could be influenced by the El Niño 2004-2005. Because the warm anomalies associated with El Niño spread from south to north, there was more evidence of this event during July 2004 in the oceanic region off the GU (present study) and in January 2005 off northern Baja [1]. El Niño 2004-2005 was not a predicted event [40] and is still under discussion for its occurrence and magnitude of surface water advection [43], but the hyperiid findings from the present and previous studies [1,3] suggest a south-to-north advection of *A. blossevillei*.

### Cross-shelf differences in species assemblages and La Niña

Hyperiid amphipods certainly present cross-shelf differences, as was established first for the California Current off Oregon [15], and further confirmed off Baja California [1]. In the present study, the influence of coastal upwelling off Punta Abreojos was consistently observed during all summers of 2002-2008 (Figs 2 and 7). This upwelling plume spread to the south in the oceanic region and, due to the topography of the area, produces an inshore-offshore split, isolating the GU, which maintains warmer temperature. The summer advance of a poleward current also increases temperatures inside the GU [21]. The upwelling front appears to be an effective barrier against the entry of amphipods from the open ocean to the coastal shelf.

The low presence of hyperiids in the GU was almost constant across all summers in the period 2002-2008, as well as in other seasons during 2005 [1]. A clear exception to this pattern was the summer of 2007, when I observed high abundance of *Simorhynchotus antennarius, Lestrigonus schizogeneios*, and *Lycaea pulex*. This could be attributed to the influence of La Niña 2007-2008 [44], promoting the biological productivity. High values of primary productivity were recorded from January to late spring 2007 in the GU [20].

However, the increase of amphipods in the GU and the offshore region during summer 2007 could also result from a seasonal effect due to delayed sampling (one month later than the rest of summers analyzed in the present study). During 2007, we are likely seeing processes more typical of autumn, when there is a seasonal increase in hyperiid abundance and diversity off Baja California [1-2]. The trend toward increasing abundance during autumn was first observed in inshore waters off Oregon for some species (*Hyperoche medusarum, Themisto pacifica*, and *Paraphronima gracilis*) but only in some years during 1963-1967 [15]. In Sagami Bay, Japan, maximuml abundance during the year occurred in September; 15 of 25 species showed this trend [31].

### Presence of gelatinous organisms enhances hyperiid abundance

The combination of seasonal and interannual effects, as well as the availability of abundant gelatinous organisms during 2007, could explain the high abundance of hyperiids in the coastal shelf of Baja California. Many amphipod species were significantly correlated with medusae and ctenophores (Table 5), which is consistent with symbiotic or parasitoid records in the literature [34,44,45]. For example, *Hyperoche medusarum* has been reported in symbiotic association with the hydromedusae *Tima Formosa* [47], *Chromatonema erythrogonon* [48], *Mitrocoma cellularia* [35], and with the ctenophores *Pleurobrachia bachei* [49], *Beroe ovata*, and *Mnemipsis leidyi* [50]. Another example is *L. schizogeneios*, reported as symbiont of hydromedusae *Clytia hemisphaerica, Liriope tetraphylla* [51], *Aequorea* sp. [45], *Leuckartiara zacae* [35] and the ctenophore *Lampea pancerina* [52].

*H. medusarum* and *L. schizogeneios*, as well as other species of the families Hyperiidae and Lestrigonidae, have many records of hyperiid-host associations, mainly involving medusa and ctenophores [45]. This is consistent with the positive correlations of *H. medusarum, L. vespuliformis*, and *L. schizogeneios* with medusae and ctenophores in the present study. *L. vespuliformis* is the only one of these three species without previous records of symbiotic associations, because it is a new species recently described [13] from the northeast Pacific near the study region. *L. vespuliformis* has been found in a few other locations, all in tropical-subtropical latitudes [1,3,14,33]. This species was considered rare [33], but in the present study it occurred frequently and showed correlations with all five gelatinous organisms. Therefore, it is highly probable that *L. vespuliformis*, which is morphologically similar to *H. medusarum*, may have a symbiotic association with one or more gelatinous species as suggested by the results of the current study.

Other abundant species in the GU during 2007 were *L. pulex* and *S. antennarius*, which both showed significant correlation with medusae and salps, and *S. antennarius* also with ctenophores. *L. pulex* has been reported in association with diverse salp species [53]. *S. antennarius* has only two symbiotic records, one with the hydromedusa *Geryonia proboscidalis* in the Mediterranean [34], and *Liriope tetraphylla* in Monterey Bay, California [46]. Both medusae species were present in the GU during the summer of 2007 (S2 Table).

## Conclusions

In conclusion, the study region is less populated with hyperiid amphipods compared to northern Baja California. The transition zone species present in northern Baja California and other northern sectors of the CCS, were also present in the GU and offshore region, but their abundances declined, particularly for *P. brevidens* and *P. gracilis*, while *S. antenarius* increased.

During summer, contrasting cross-shelf differences in species assemblages were observed with high interannual variability. Active upwelling off Punta Abreojos forms a plume separating the inshore and offshore regions, preventing entry of amphipods to the coastal shelf. Changes in temperature and the proliferation of gelatinous organisms during La Niña 2007-2008 promoted the occupancy of the GU by hyperiids. Future studies in other seasons will allow us to corroborate the spatial-temporal tendencies observed in the present study, and to further investigate whether there are similar progressions of amphipods on the coastal shelf.

## Acknowledgements

In memory of Bill Peterson, colleague in the study of the California Current and zooplankton ecology. Thanks to the people participating in IMECOCAL cruises and laboratory assistance. Thanks to Laura E. Lilly for criticism of the manuscript. Financial support provided by CONACYT grants (129611, 23804, 23947), and the CICESE project *Variability of zooplankton in function of climatic changes of diverse scale*.

## Supporting information

**S1 Table. Oceanographic stations**. Location and sampling date of oceanographic **s**tations during the IMECOCAL cruises at the Gulf of Ulloa and offshore region.

**S2 Table. Hyperiid species list**. Hyperiid species separated by frequency of occurrence (FO) in dominant (31-80%), common (10-30%), and rare (<10%) in 108 samples analyzed. The Geometric Mean (GM) and the mean are also shown, combining data of samples collected during nighttime (N = 88). (*) indicate species found in daytime samples only.

**S3 Table. Medusae abundance**. Abundance of two species of limnomedusae during the summer of 2007 (IMECOCAL cruise 0708).

**S1 Fig. Summertime abundance of selected species from the Gulf of Ulloa, Baja California**.**- Part 1**. Mean (± 95% confidence interval) in the offshore and onshore regions for species in the infraorders Physososomata (a) and Physocephalata (b-f): families Scinidae (a), Paraphronimidae (b), Vibilidae (c), and Phrosinidae (d-f).

**S2 Fig. Summertime abundance of selected species from the Gulf of Ulloa, Baja California**.**- Part 2**. Mean (± 95% confidence interval) in the offshore and onshore regions for species in the infraorder Physocephalata: families Hyperiidae (a-b), Lestrigonidae (c-d), and Phronimidae (e-f).

**S3 Fig. Summertime abundance of selected species from the Gulf of Ulloa, Baja California**.**- Part 3**. Mean (± 95% confidence interval) in the offshore and onshore regions for species in the infraorder Physocephalata: families Eupronoidae (a-b), Platyscelidae (c-d), and Lycaeidae (e).

